# Similar efficacies of selection shape mitochondrial and nuclear genes in *Drosophila melanogaster* and *Homo sapiens*

**DOI:** 10.1101/010355

**Authors:** Brandon S. Cooper, Chad R. Burrus, Chao Ji, Matthew W. Hahn, Kristi L. Montooth

## Abstract

Deleterious mutations contribute to polymorphism even when selection effectively prevents their fixation. The efficacy of selection in removing deleterious mitochondrial mutations from populations depends on the effective population size (*N*_*e*_) of the mtDNA, and the degree to which a lack of recombination magnifies the effects of linked selection. Using complete mitochondrial genomes from *Drosophila melanogaster* and nuclear data available from the same samples, we re-examine the hypothesis that non-recombining animal mtDNA harbor an excess of deleterious polymorphisms relative to the nuclear genome. We find no evidence of recombination in the mitochondrial genome, and the much-reduced level of mitochondrial synonymous polymorphism relative to nuclear genes is consistent with a reduction in *N*_*e*_. Nevertheless, we find that the neutrality index (*NI*), a measure of the excess of nonsynonymous polymorphism relative to the neutral expectation, is not significantly different between mitochondrial and nuclear loci. Reanalysis of published data from *Homo sapiens* reveals the same lack of a difference between the two genomes, though small samples in previous studies had suggested a strong difference in both species. Thus, despite a smaller *N*_*e*_, mitochondrial loci of both flies and humans appear to experience similar efficacies of selection as do loci in the recombining nuclear genome.

## INTRODUCTION

The effective size of a population (*N*_*e*_) impacts how effectively selection removes deleterious mutations and fixes advantageous mutations. The unique genetics of the mitochondrial genome (mtDNA) are thought to reduce its *N*_*e*_ relative to the nuclear genome, via haploid, uniparental inheritance, the mitochondrial bottleneck in the maternal germline, and a lack of recombination that decreases *N*_*e*_ via selection on linked sites (Hill and Robertson 1966; Maynard Smith and Haigh 1974; Gillespie 2000; Meiklejohn *et al.* 2007; White *et al.* 2008; Charlesworth 2012). In addition, cytoplasmic transmission can link the mtDNA to selfish cytoplasmic elements (e.g., *Wolbachia* in insects) that may sweep through populations, further decreasing mitochondrial *N*_*e*_ and possibly increasing mitochondrial substitution rates via the fixation of slightly deleterious mutations (Shoemaker *et al.* 2004). For these reasons it has been widely hypothesized that selection is less effective in mitochondrial genomes than in their nuclear counterparts, and that mitochondrial genomes may accumulate greater numbers of deleterious substitutions (Lynch 1996; Lynch 1997). Analyses of sequence data in *Drosophila* and mammals have largely supported the conclusion that mtDNA harbor significant levels of slightly deleterious polymorphism (Ballard and Kreitman 1994; Rand and Kann 1996; Nachman 1998; Rand and Kann 1998; Weinreich and Rand 2000).

*N*_*e*_ is not the only evolutionary parameter that distinguishes mitochondrial and nuclear genomes. The distinct functional landscape of the mitochondrial genome likely affects the distribution of selective effects (*s*) of mutations that arise in this genome. Animal mitochondrial genomes typically encode regulatory information for replication and transcription nested within a hypervariable region (also known as the D-loop, control, or A+T-rich region), 22 tRNAs, 2 ribosomal components, and 13 protein-coding genes—all core components of oxidative phosphorylation (OXPHOS). Outside of the control region, there is little noncoding DNA in animal mtDNAs. In Drosophilids, 99% of the genome outside of the control region encodes DNA and RNA genes with highly conserved sequences that function in mitochondrial protein synthesis and aerobic respiration (Clary and Wolstenholme 1985; Wolstenholme and Clary 1985; Ballard 2000; Montooth *et al.* 2009). This suggests that the distribution of selective effects in the mtDNA may be shifted towards larger negative effects on fitness.

The mutational landscape of the mtDNA also differs from the nuclear genome. In most animal taxa the mitochondrial mutation rate greatly exceeds that of the nuclear genome (Lynch *et al.* 2008), and the mitochondrial mutational process is also highly biased (Montooth and Rand 2008). For example, nearly all mitochondrial mutations in *D. melanogaster* change a G:C base pair to an A:T (Haag-Liautard *et al.* 2008). When combined with the strong A+T-bias in this mitochondrial genome, where 95% of third codon positions are an A or a T (Montooth *et al.* 2009), this indicates that the most commonly occurring mutations in protein-coding loci of the *Drosophila* mtDNA will change an amino acid. Relative to the nuclear genome, animal mitochondrial genomes thus experience a greater mutational pressure that can also be biased in some taxa towards nonsynonymous mutations; these are likely to have deleterious effects in a molecule that encodes such highly conserved functions.

Some of the strongest population genetic patterns in support of distinct selective pressures acting on mitochondrial and nuclear genomes come from analyses of the neutrality index (*NI*) (Rand and Kann 1996; Nachman 1998; Rand and Kann 1998; Weinreich and Rand 2000). *NI* is a summary statistic of the deviation from the neutral expectation in the McDonald-Kreitman test (Mcdonald and Kreitman 1991; Rand and Kann 1996), and is calculated from counts of synonymous and nonsynonymous polymorphic and fixed sites within and between related species. Weakly deleterious nonsynonymous mutations that segregate in the population but that will not contribute to divergence lead to a value of *NI* greater than 1. When the efficacy of selection is decreased, the expectation is that the number of segregating weakly deleterious polymorphisms will increase; this is the pattern that has been observed in mtDNA. Meta-analyses of McDonald-Kreitman tables and their associated *NI* values for mitochondrial and nuclear loci in animals have concluded that *NI* is predominantly greater than the neutral expectation of 1 for the mtDNA (Rand and Kann 1996; Nachman 1998; Rand and Kann 1998; Betancourt *et al.* 2012) and exceeds the average *NI* of the nuclear genome (Weinreich and Rand 2000). Although the relative sparseness of the data was recognized early on (Nachman 1998), and the conclusions were largely limited to how selection shapes animal mtDNA, these patterns are often taken as evidence that selection is largely ineffective in the mtDNA because of its reduced *N*_*e*_, and that mitochondrial genomes are expected to harbor more deleterious polymorphisms than do their nuclear counterparts.

Here we revisit this pattern using new, complete mitochondrial genomes from *D. melanogaster* that we compare with published nuclear data from the same samples (Langley *et al.* 2012) and with available human data from both nuclear and mitochondrial genomes (Bustamante *et al.* 2005; Just *et al.* 2008; Rubino *et al.* 2012). We find little evidence that the effects of selection differ on average between mitochondrial and nuclear genomes of flies or humans, despite evidence that there is much reduced *N*_*e*_ due to a lack of recombination and linkage with the cytoplasm. We discuss reasons why *NI* is, on average, similar between mitochondrial and nuclear loci, despite the distinct population genetic properties of these two genomes.

## MATERIALS AND METHODS

### *D. melanogaster* mtDNA assembly, annotation, and estimates of sequence diversity

Raw sequence read files from 38 genetic lines of *D. melanogaster* from Raleigh, NC, USA (Mackay *et al.* 2012) sequenced by the 50 Genomes subproject of the *Drosophila* Population Genomics Project (DPGP) (Langley *et al.* 2012) were downloaded from the NCBI Sequence Read Archive. We used the Burrows-Wheeler Aligner (BWA), and specifically the fast and accurate short read alignment with Burrows-Wheeler Transform (Li and Durbin 2009), to map sequence reads to the *D. melanogaster* mitochondrial reference genome (NC_001709). We allowed up to 5 gaps, 5 gap extensions, and 5 mismatches per aligned read, but few reads needed such flexibility and most were filtered out in later steps. Using SAMtools, we post-processed the alignments to filter out low-quality alignments and to detect single nucleotide polymorphisms (SNPs) (Li *et al.* 2009). SNPs with a quality score greater than 20 and indels with a quality score greater than 50 were kept for further analyses. We then generated a consensus sequence for each of the *D. melanogaster* mtDNAs listed in Table S1. Due to high variance in coverage across the A+T-rich D-loop, we did not include this region in our final assemblies or analyses.

We annotated the consensus sequence for each mtDNA using the GenBank annotation of the *D. melanogaster* reference sequence (NC_001709). Using ClustalW (Larkin *et al.* 2007), we performed a whole genome alignment, as well as gene-specific alignments, of each consensus sequence to the reference sequence and to the outgroup species *D. yakuba* (NC_001322). There are very few indels in the coding regions of *Drosophila* mtDNA (Montooth *et al.* 2009), making alignment straightforward. From these alignments we calculated expected heterozygosity (π), the number of segregating sites (*S*), and Watterson’s *θ*_*W*_ (Watterson 1975) as measures of sequence diversity. The mitochondrial haplotype network was inferred from 80 segregating sites in the coding region of the mtDNA for which there were no missing or ambiguous data using TCS version 1.21 (Clement *et al.* 2000).

### Tests for recombination in the *D. melanogaster* mtDNA

We estimated linkage disequilibrium (LD) between all pairs of mitochondrial SNPs using the statistic *D’* (Lewontin 1964), where *D*’ = 0 indicates no LD and ∣*D*’∣ = 1 indicates perfect LD. Because recombination erodes LD as a function of distance, a negative correlation between ∣*D*’∣ and genetic distance between pairs of SNPs has been used as evidence for recombination (Awadalla *et al.* 1999). To test this prediction we looked for significant negative correlations between ∣*D*’∣ and genetic distance. We also conducted these same tests using another statistical measure of association, *r*^2^ (Hill and Robertson 1966), which is more robust to variation in mutation rates (Awadalla *et al.* 1999; Meunier and Eyre-Walker 2001; Innan and Nordborg 2002). We calculated these correlations using a variety of minor allele frequency cutoffs (Table S2). We also tested for the presence of all four genotypes at pairs of SNPs (the “four-gamete test”; Hudson and Kaplan 1985) using DNAsp version 5 (Rozas *et al.* 2003).

### Neutrality tests

Using π and *θ*_*W*_, we calculated Tajima’s *D* (Tajima 1989), which is expected to be 0 for a neutrally evolving locus. Demographic effects will skew the site-frequency spectrum of both synonymous and nonsynonymous polymorphisms at a locus. Contrasting Tajima’s *D* between nonsynonymous and synonymous polymorphisms therefore tests whether nonsynonymous alleles experience a greater skew in frequency relative to putatively neutral synonymous alleles, indicative of selection (Rand and Kann 1996). We implemented this analysis using the heterogeneity test (Hahn *et al.* 2002), which simulates 10,000 genealogies with no recombination using the values of synonymous and nonsynonymous *S* calculated from the data and compares the estimated difference in Tajima’s *D* to the random distribution of differences between synonymous and nonsynonymous polymorphisms. We calculated several other summaries of the site-frequency spectrum, including Fu and Li’s *D*, which characterizes the proportion of mutations on external and internal branches of a genealogy (Fu and Li 1993) and Fay and Wu’s *H*, which tests for an excess of high-frequency derived alleles in a sample relative to the neutral expectation (Fay and Wu 2000). These latter statistics were calculated using a set of 80 segregating sites in the coding region of the mtDNA for which there were no missing or ambiguous data. Significance was determined using 10,000 coalescent simulations as implemented in DNAsp version 5 (Rozas *et al.* 2003).

We conducted McDonald-Kreitman (MK) tests (Mcdonald and Kreitman 1991) using Fisher’s exact tests of the two-by-two contingency tables of counts of nonsynonymous and synonymous polymorphisms (*P*_*N*_ and *P*_*S*_) within *D. melanogaster* and nonsynonymous and synonymous fixed differences (*D*_*N*_ and *D*_*S*_) between *D. melanogaster* and *D. yakuba*. Polymorphic sites within *D. melanogaster* only contributed to fixed differences if the allele in *D. yakuba* was not present in *D. melanogaster.* For codons with more than one change, we calculated the number of nonsynonymous and synonymous differences as the average over all possible mutational pathways between codons (Nei and Gojobori 1986). For any gene with a count of 0 in any cell of the MK table, we added a count of 1 to all cells (Sheldahl *et al.* 2003; Presgraves 2005). 23% of thirteen *D. melanogaster* mitochondrial genes, 9.5% of 6,113 *D. melanogaster* nuclear genes, 0% of thirteen *H. sapiens* mitochondrial genes, and 73% of 11,624 *H. sapiens* nuclear genes required these additional counts. Fisher’s exact tests were performed using R version 2.15.1 (R Core Team 2012).

We calculated *NI*—the ratio of *P*_*N*_*/P*_*S*_ to *D*_*N*_*/D*_*S*_—as a summary statistic of the MK table (Rand and Kann 1996). Assuming that selection is constant, the neutral expectation is that *D*_*N*_*/D*_*s*_ will equal *P*_*N*_*/P*_*S*_ (Kimura 1983; Mcdonald and Kreitman 1991), and *NI* is expected to be 1. When the MK test is significant, an *NI* less than 1 indicates a significant excess of nonsynonymous fixed differences and an *NI* greater than 1 indicates a significant excess of nonsynonymous polymorphisms. We calculated several modified statistics similar to *NI*, including 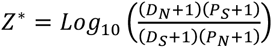 (Presgraves 2005) and the direction of selection 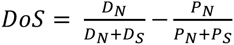 which is an unbiased summary statistic of the two-by-two MK table (Stoletzki and Eyre-Walker 2011). The sign of these latter two statistics is more intuitive; negative values are consistent with an excess of weakly deleterious (negatively selected) polymorphisms and positive vales are consistent with an excess of advantageous (positively selected) substitutions.

The short length and reduced polymorphism of mitochondrial genes decreases the power of the MK test and upwardly biases *NI* (Weinreich and Rand 2000). Because of this, we also performed MK tests and calculated *NI* using the summed counts of polymorphic and divergent sites for each of the OXPHOS complexes: Complex I (NADH dehydrogenase, ND), Complex IV (cytochrome *c* oxidase, CO) and Complex V (ATP synthetase, ATPase). Cytochrome B (CytB) is the only Complex III gene encoded by the mtDNA. Stoletzki and Eyre-Walker (2011) emphasize that contingency data should not generally be summed, particularly when there is heterogeneity among contingency tables, and they provide an unbiased estimator of overall *NI* for combining counts, 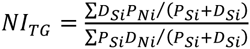 We calculated this statistic and used the DoFE software package (Stoletzki and Eyre-Walker 2011) to calculate bootstrap confidence intervals and to conduct Woolf’s tests of homogeneity (Woolf 1955). The only data set with significant heterogeneity was the *D. melanogaster* nuclear gene set (*P<*0.0001). The same approaches were used to analyze polymorphism and divergence in the human data sets (see below), as well as in a subset of the mitochondrial haplotypes reported in our study that were independently sequenced and assembled by Richardson et al. (2012) (Table S3).

### Comparisons of mitochondrial and nuclear *NI* in flies and humans

To compare patterns of polymorphism and divergence between mitochondrial and nuclear genomes, we obtained existing data for nuclear genes in *D. melanogaster* and for nuclear and mitochondrial genes in *Homo sapiens.* Counts of polymorphic and divergent sites for *D. melanogaster* nuclear genes were obtained from the DPGP analysis of the same 38 genomes from Raleigh, NC, USA, using *D. yakuba* as the outgroup species (Langley *et al.* 2012). The human nuclear data from Bustamante et al. (2005) included counts of polymorphic and divergent sites from 19 African Americans and 20 European Americans, using the chimpanzee *Pan troglodytes* as an outgroup species. We calculated the number of polymorphic and divergent sites for human mitochondrial genes using mtDNA sequences from 19 African Americans (Just *et al.* 2008), 20 European Americans (Rubino *et al.* 2012), and the chimpanzee mitochondrial reference genome D38113.1 (Horai *et al.* 1995). We also analyzed subsets of the human mitochondrial sequence data to characterize the sensitivity of *NI* to sampling (Table S4). Table S5 provides accession numbers for the human mtDNA sequences used. Comparisons of the distributions of *NI* between data sets were performed using Mann-Whitney *U* tests in R version 2.15.1 (R Core Team 2012).

## RESULTS

### An excess of low- and high-frequency derived mitochondrial polymorphisms

We assembled 14,916 bp of sequence containing the transcribed regions of the mtDNA with a median coverage of 32x for 38 genetic lines sampled from a single population of *D. melanogaster* in Raleigh, NC (Langley *et al.* 2012; Mackay *et al.* 2012)(Table S1). Over 98% of these nucleotides encode the 13 protein-coding genes, 22 tRNAs and two ribosomal RNAs. The per-site expected heterozygosity in this region (π) of the mtDNA was 0.0008. We identified 137 segregating sites in this population sample, 103 of which were in protein-coding genes. Median heterozygosity in protein coding genes was 0.0023 per synonymous site and 0.0002 per nonsynonymous site. Silent site heterozygosity was significantly lower in mitochondrial genes relative to nuclear genes (Mann-Whitney *U*, mtDNA v. X chromosome, *P*_*MWU*_ = 0.00002; mtDNA v. autosomes, *P*_*MWU*_ < 0.00001) and was only 0.16 times that of the autosomes (Figure 1A), lower than what is expected if the mtDNA has an effective population size that is one-quarter that of the autosomes.

**Figure 1.**
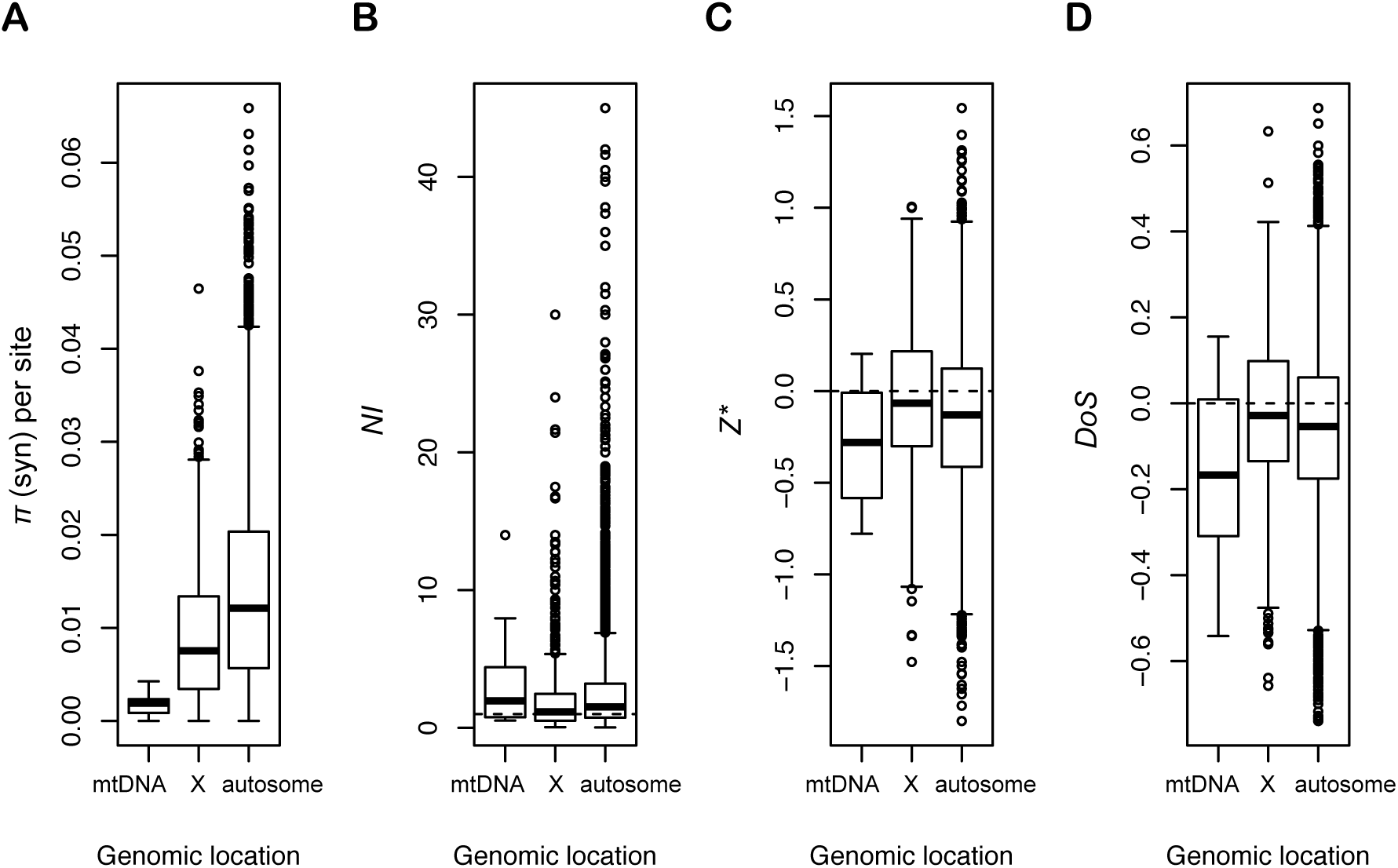
Effect of genomic location on silent-site heterozygosity and on summary statistics of polymorphism and divergence in *D. melanogaster.* (A) Genomic location has a significant effect on silent-site heterozygosity (*P*_*MWU*_ < 0.001 for all pairwise contrasts), consistent with predicted differences in *N*_*e*_ between these chromosomes. The ratio of median mitochondrial to autosomal silent site heterozygosity was 0.157, less than predicted for neutral sites if mitochondrial *N*_*e*_ is one quarter that of the autosomes. mtDNA, X-chromosome, and autosome data sets contained 12, 1255, and 8073 genes, respectively. (BD) Distributions of neutrality indices *NI*, *Z\** and *DoS* are similar between mitochondrial and nuclear genes despite differences in *N*_*e*_. Dashed lines represent the neutral expectations for these statistics. Three nuclear loci for which *NI* exceeded 50 were excluded from (B) to improve visualization. See main text for description of statistics, number of genes in each dataset and statistical results.

In addition to there being very few segregating sites in the *D. melanogaster* mtDNA, polymorphisms at these sites were skewed towards low frequencies (Figure 2A,B), as evidenced by consistently negative values of Tajima’s *D* (Table 1). Tajima’s *D* across the mtDNA was -2.607 and differed significantly from the neutral expectation of 0 (*S*=80 for coding sites with no missing data, *P<*0.0001), as did Fu and Li’s *D* (*D* = -2.67, *P<*0.05). The minor allele frequency for unpolarized synonymous polymorphisms was always less than 11%, and all but one of the nonsynonymous polymorphisms was a singleton (Figure 2A,B). Using *D. yakuba* as an outgroup revealed that the derived allele was nearly fixed for 44% of segregating synonymous sites, while there was only a single derived nonsynonymous polymorphism at high frequency (Figure 2C,D). Thus, the mitochondrial genome was essentially devoid of intermediate frequency polymorphisms, with derived synonymous mutations at either very high (greater than 89%) or very low (less than 11%) frequencies and nearly all derived nonsynonymous polymorphisms at frequencies less than 5%. This skew towards high-frequency derived alleles resulted in a significant negative value of Fay and Wu’s *H* statistic (*H* = -41.2, *P* = 0.005).

**Table 1.** Synonymous and nonsynonymous variation in *D. melanogaster* mitochondrial genes and OXPHOS complexes.

| Gene/Complex<sup>a</sup> | Synonymous sites<sup>b</sup> |  |  |  |  | Nonsynonymous sites |  |  |  |  |
| --- | --- | --- | --- | --- | --- | --- | --- | --- | --- | --- |
| | n (bp) | S | $\pi$ | $\Theta_W$ | $D$ | n (bp) | S | $\pi$ | $\Theta_W$ | $D$ |
| *COI* | 347 | 16 | 1.57 | 3.81 | -1.91 | 1183 | 0 | 0 | 0 | undef |
| *COII* | 142 | 5 | 0.31 | 1.19 | -1.90 | 539 | 1 | 0.05 | 0.24 | -1.13 |
| *COIII* | 169.33 | 9 | 0.72 | 2.14 | -1.96 | 613.67 | 1 | 0.05 | 0.24 | -1.13 |
| *NDI* | 194 | 3 | 0.16 | 0.71 | -1.72 | 739 | 3 | 0.16 | 0.71 | -1.72 |
| *NDII* | 198 | 5 | 0.41 | 1.19 | -1.69 | 822 | 6 | 0.34 | 1.43 | -2.07 |
| *NDIII* | 68 | 1 | 0.05 | 0.24 | -1.12 | 280 | 1 | 0.05 | 0.24 | -1.12 |
| *NDIV* | 273.33 | 7 | 0.51 | 1.67 | -1.95 | 1061.67 | 7 | 0.38 | 1.67 | -2.17 |
| *NDIVL* | 56 | 1 | 0.05 | 0.24 | -1.13 | 229 | 2 | 0.11 | 0.48 | -1.49 |
| *NDV* | 351 | 10 | 0.68 | 2.38 | -2.16 | 1365 | 4 | 0.21 | 0.95 | -1.88 |
| *NDVI* | 103.33 | 4 | 0.26 | 0.95 | -1.75 | 415.67 | 2 | 0.11 | 0.48 | -1.49 |
| Complex I | 1243.66 | 31 | 2.12 | 7.38 | -2.48 | 4912.34 | 25 | 1.35 | 5.95 | -2.64 |
| Complex III | 239 | 5 | 0.55 | 1.19 | -1.38 | 892 | 2 | 0.16 | 0.48 | -1.29 |
| Complex IV | 658.33 | 30 | 2.60 | 7.14 | -2.21 | 2335.67 | 2 | 0.11 | 0.48 | -1.49 |
| Complex V | 174.67 | 2 | 0.16 | 0.71 | -1.72 | 650.33 | 6 | 0.32 | 1.43 | -2.10 |
| Total | 2315.66 | 68 | 5.43 | 16.42 | -2.44 | 8790.34 | 35 | 1.93 | 8.33 | -2.70 |

<sup>a</sup> Complex I, ND, 7 loci; Complex III, Cyt-b, 1 locus; Complex IV, CO, 3 loci; Complex V, ATPase, 2 overlapping loci
<sup>b</sup> For synonymous and nonsynonymous sites we calculated the number of segregating sites ( $S$ ), heterozygosity ( $\pi$ ), Watterson's $\Theta_W$ , and Tajima's $D$ . The heterogeneity test for differences between synonymous and nonsynonymous $D$ was never significant ( $P > 0.35$ for all comparisons).

**Figure 2.**
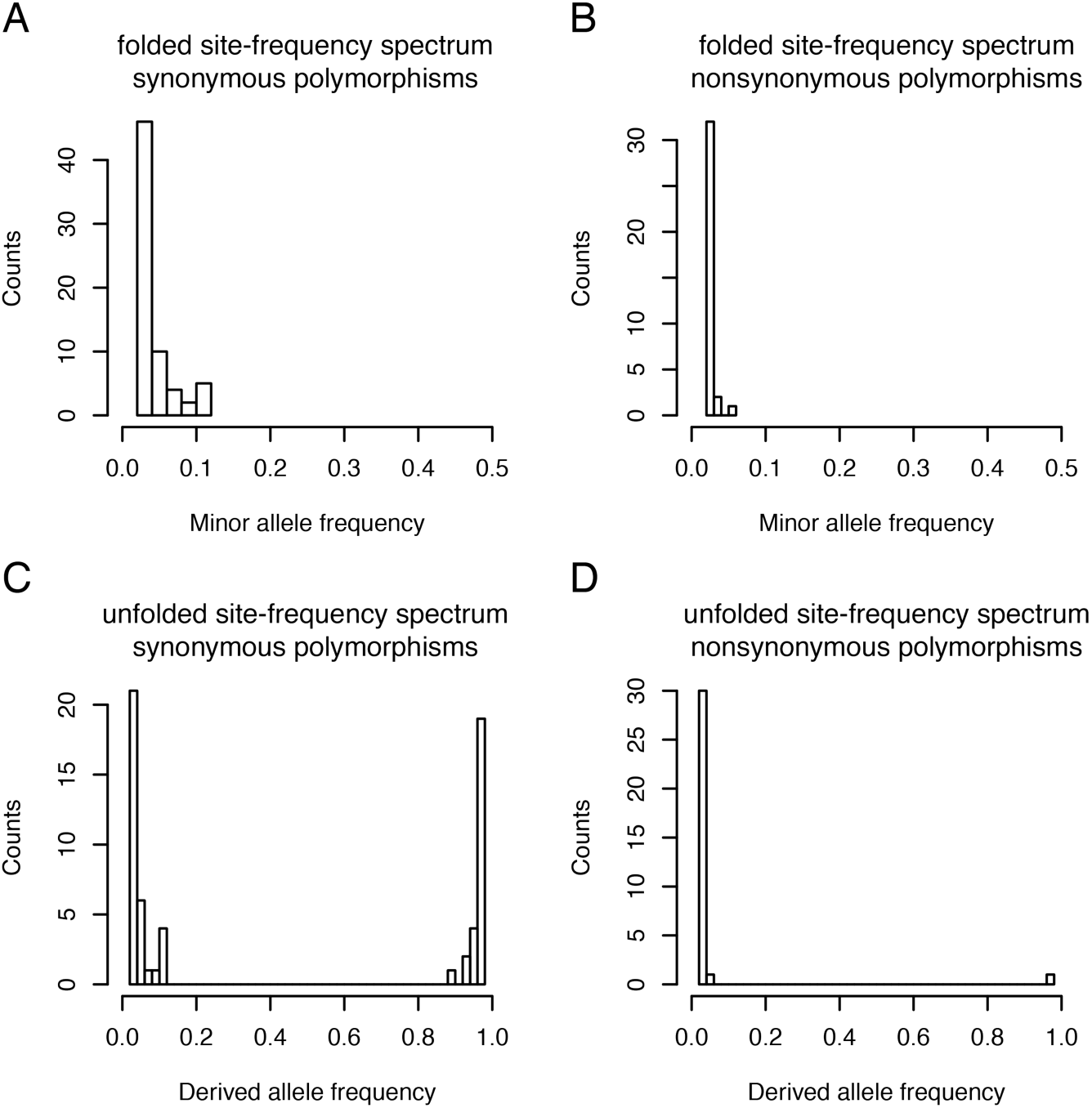
Site-frequency spectra of synonymous and nonsynonymous polymorphisms in the *D. melanogaster* mtDNA. (A, B) Folded site-frequency spectra for synonymous and nonsynonymous segregating sites across the mitochondrial coding region reveal that mitochondrial polymorphisms are skewed to low frequencies and are more so for nonsynonymous sites. (C, D) Unfolded site-frequency spectra reveal that derived, synonymous polymorphisms are almost equally likely to be at low frequency (56% of 59 sites at frequencies less than 0.11) or nearly fixed (44% of 59 sites at frequencies greater than 0.89), while derived, nonsynonymous polymorphisms are nearly always present as singletons (94% of 32 sites). There are essentially no mitochondrial polymorphisms at intermediate frequencies. Sites were omitted from the unfolded site frequency spectra if neither allelic state was shared with *D. yakuba*. The number of sites included in each distribution is 67 (A), 35 (B), 59 (C), and 32 (D).

### A partial selective sweep in the *D. melanogaster* mtDNA

The large fraction of derived alleles at high frequencies is a consequence of the haplotype structure of this sample (Figure 3). Nearly 30% of individuals in this population shared an identical mitochondrial haplotype and an additional 66% of individuals differed from this haplotype by only one to five mutations. The two remaining haplotypes (RAL-639 and RAL-335) were highly divergent from this common haplotype group, contributing nearly half of the segregating sites to the population sample. These two haplotypes shared the ancestral state with *D. yakuba* at 17 of the 23 derived high-frequency synonymous polymorphisms (i.e., they have the low-frequency ancestral allele; Figure 2C). When these two haplotypes were removed from the analysis, there remained a strong skew toward rare alleles (Tajima’s *D* = -2.31, *P* < 0.01; Fu and Li’s *D* = -3.14, *P* < 0.02), but Fay and Wu’s *H*, which is sensitive to the number of high-frequency derived alleles, was only weakly significant (*H* = -10.34, *P* = 0.043). The remaining six derived, high-frequency synonymous polymorphisms, as well as the single derived, high-frequency nonsynonymous polymorphism, were the result of single mtDNAs within the common haplotype group having the same allelic state as *D. yakuba*. Given the lack of recombination in the mtDNA (see below), these are likely new, rather than ancestral, mutations. Six of these seven mutations would have changed a C or G to a T or A, consistent with the mutation bias in the *D. melanogaster* mtDNA (Haag-Liautard *et al.* 2008).

**Figure 3.**
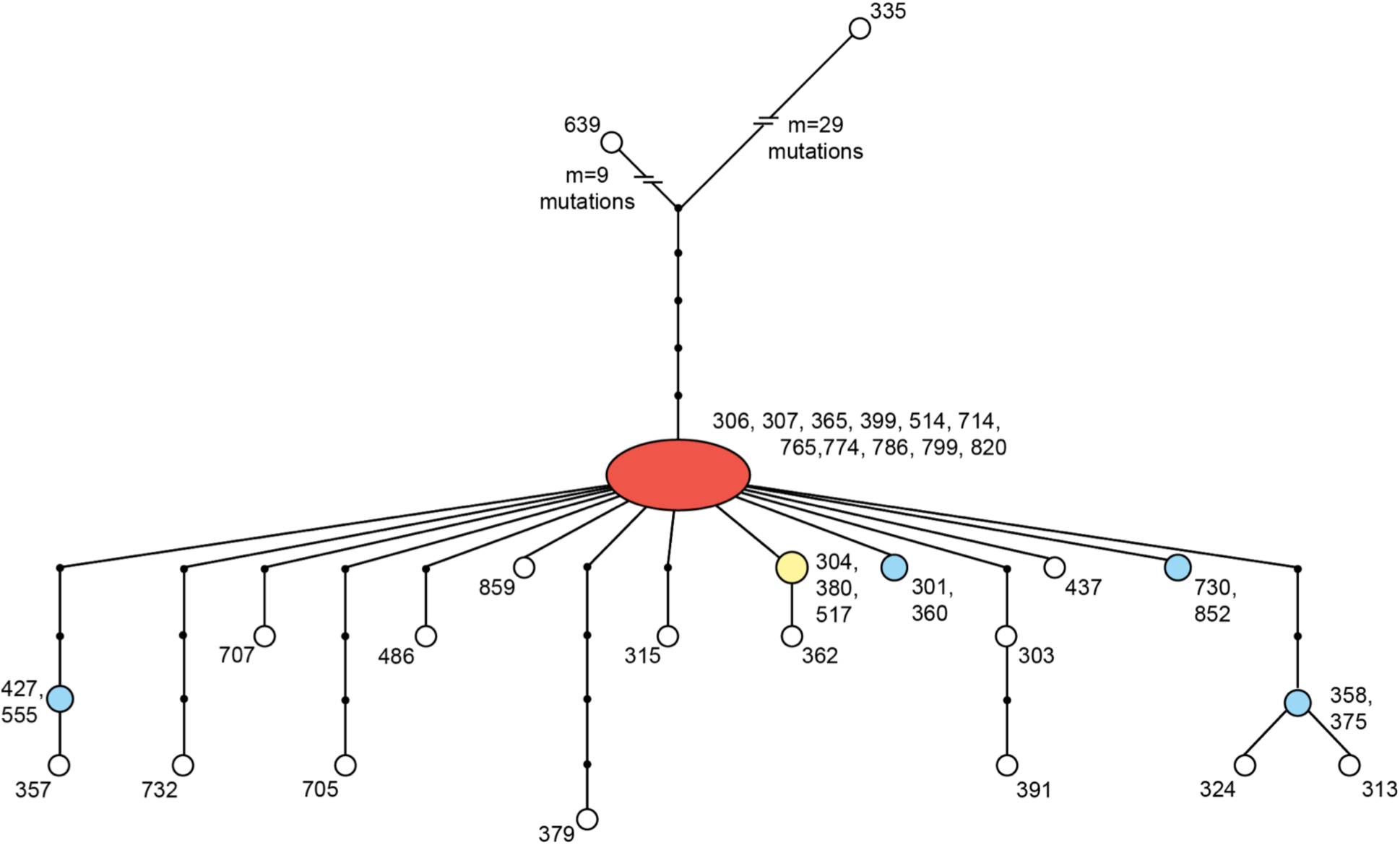
Haplotype network for 38 *D. melanogaster* mtDNAs sampled from Raleigh, NC. The network, inferred from 80 coding region SNPs with no missing information, reveals that nearly 30% of individuals sampled (11/38) share the same common haplotype (red) and an additional 65% of individuals carry a haplotype only a few mutations away from this haplotype. This common set of mitochondrial haplotypes is highly diverged from the two other mtDNAs sampled in the population; lines RAL-639 and RAL-335 differ from the common haplotype at 14 and 34 SNPs, respectively. At least one of these two haplotypes carries the ancestral state (shared with *D. yakuba*) at 38% of these SNPs. Numbers represent the Raleigh (DGRP) line carrying the haplotype. Red, yellow, blue and white nodes were present in 11, 3, 2 and 1 lines, respectively.

### No evidence for recombination in the *D. melanogaster* mtDNA

We tested for a negative correlation between linkage disequilibrium (LD) and the distance between each pair of polymorphic sites in the *D. melanogaster* mitochondrial genome, as a signature of the decay of LD over distance via recombination (Awadalla *et al.* 1999). There was no evidence for a decrease in LD with increasing distance between sites, regardless of the measure of LD or the minor allele cutoff used (Table S2). There were no pairs of polymorphic sites for which all four gametes were present (Hudson and Kaplan 1985; Bruen *et al.* 2006), further supporting an absence of effective recombination.

### Weakly deleterious polymorphism in the *D. melanogaster* mtDNA

The skew in the site-frequency spectrum toward rare alleles (Figure 2) resulted in negative values of Tajima’s *D* across the entire mtDNA (Table 1). There was no evidence that the skew toward rare alleles differed between synonymous and nonsynonymous polymorphisms (Figure 2A,B): heterogeneity tests (Hahn *et al.* 2002) of Tajima’s *D* between synonymous and nonsynonymous sites were never significant (*P* > 0.35 for all genes and complexes). However, unfolding the frequency spectra revealed that the large number of high-frequency derived sites were nearly all synonymous (Figure 2C,D). This would indicate that the haplotype that has increased in frequency carried many more synonymous than nonsynonymous polymorphisms. Given that the mutation bias in the *D. melanogaster* mtDNA greatly favors nonsynonymous mutations (Haag-Liautard *et al.* 2008), this pattern suggests a history of effective purifying selection removing mitochondrial haplotypes that contain nonsynonymous polymorphisms. Furthermore, all nonsynonymous polymorphisms that have arisen on the common mitochondrial haplotype are present at very low frequencies.

The current distribution of polymorphisms relative to divergence showed little evidence for an excess of segregating deleterious polymorphisms. No single gene and only one OXPHOS complex significantly departed from the neutral expectation after Bonferroni correction (Table 2 and Table 3); the MK test was significant for Complex V (ATPase), with an excess of nonsynonymous polymorphism (Fisher’s exact test, *P*_*FET*_ = 0.007). For the entire set of protein-coding mitochondrial genes, there was a slight excess of nonsynonymous polymorphism relative to the neutral expectation that was not significant after Bonferroni correction (*P*_*FET*_ = 0.041). Using the unbiased estimator *NI*_*TG*_ (Stoletzki and Eyre-Walker 2011) did not change this inference: *NI*_*TG*_ for mitochondrial-encoded proteins was 1.67, but the confidence intervals on *NI*_*TG*_ nearly included the neutral expectation of 1 (Table 3).

**Table 2.** Counts of polymorphic (*P*) and divergent (*D*) nonsynonymous (*N*) and synonymous (*S*) sites along with summary statistics of the McDonald-Kreitman table for *D. melanogaster* mitochondrial genes.

| Gene | $P_N$ | $P_S$ | $D_N$ | $D_S$ | $NI^a$ | $Z^{*b}$ | $DoS^c$ |
| --- | --- | --- | --- | --- | --- | --- | --- |
| *ATPase6* | 5 | 2 | 11 | 35 | 7.955 | -0.778 | -0.475 |
| *ATPase8* | 1 | 0 | 2 | 8 | 6 | -0.778 | undef |
| *COI* | 0 | 16 | 8 | 101 | 0.667 | 0.176 | undef |
| *COII* | 1 | 5 | 6 | 39 | 1.3 | -0.280 | -0.033 |
| *COIII* | 1 | 9 | 8.5 | 47.5 | 0.621 | -0.009 | 0.052 |
| *CytB* | 2 | 5 | 17.5 | 67.5 | 1.543 | -0.267 | -0.080 |
| *NDI* | 3 | 3 | 11 | 45 | 4.091 | -0.584 | -0.304 |
| *NDII* | 6 | 5 | 25 | 41 | 1.968 | -0.275 | -0.167 |
| *NDIII* | 1 | 1 | 5 | 22 | 4.4 | -0.584 | -0.315 |
| *NDIV* | 7 | 7 | 24 | 63 | 2.625 | -0.408 | -0.224 |
| *NDIVL* | 2 | 1 | 1 | 7 | 14 | -0.778 | -0.542 |
| *NDV* | 4 | 10 | 55.833 | 107.167 | 0.768 | 0.063 | 0.057 |
| *NDVI* | 2 | 4 | 21.5 | 22.5 | 0.523 | 0.203 | 0.155 |

<sup>a</sup> For *ATPase8* and *COI*, a count of 1 was added to each cell when calculating $NI = \frac{D_S P_N}{D_N P_S}$ . For no gene was the neutral expectation rejected via a Fisher's exact test of the MK table at the table-wise Bonferroni corrected $P=0.004$ .
<sup>b</sup> $Z^* = \log_{10} \left( \frac{(D_N+1)(P_S+1)}{(D_S+1)(P_N+1)} \right)$ , as in (Presgraves 2005).
<sup>c</sup> $DoS = \frac{D_N}{D_N+D_S} - \frac{P_N}{P_N+P_S}$ was not calculated for loci with a count of zero (Stoletzki and Eyre-Walker 2011).

**Table 3.**
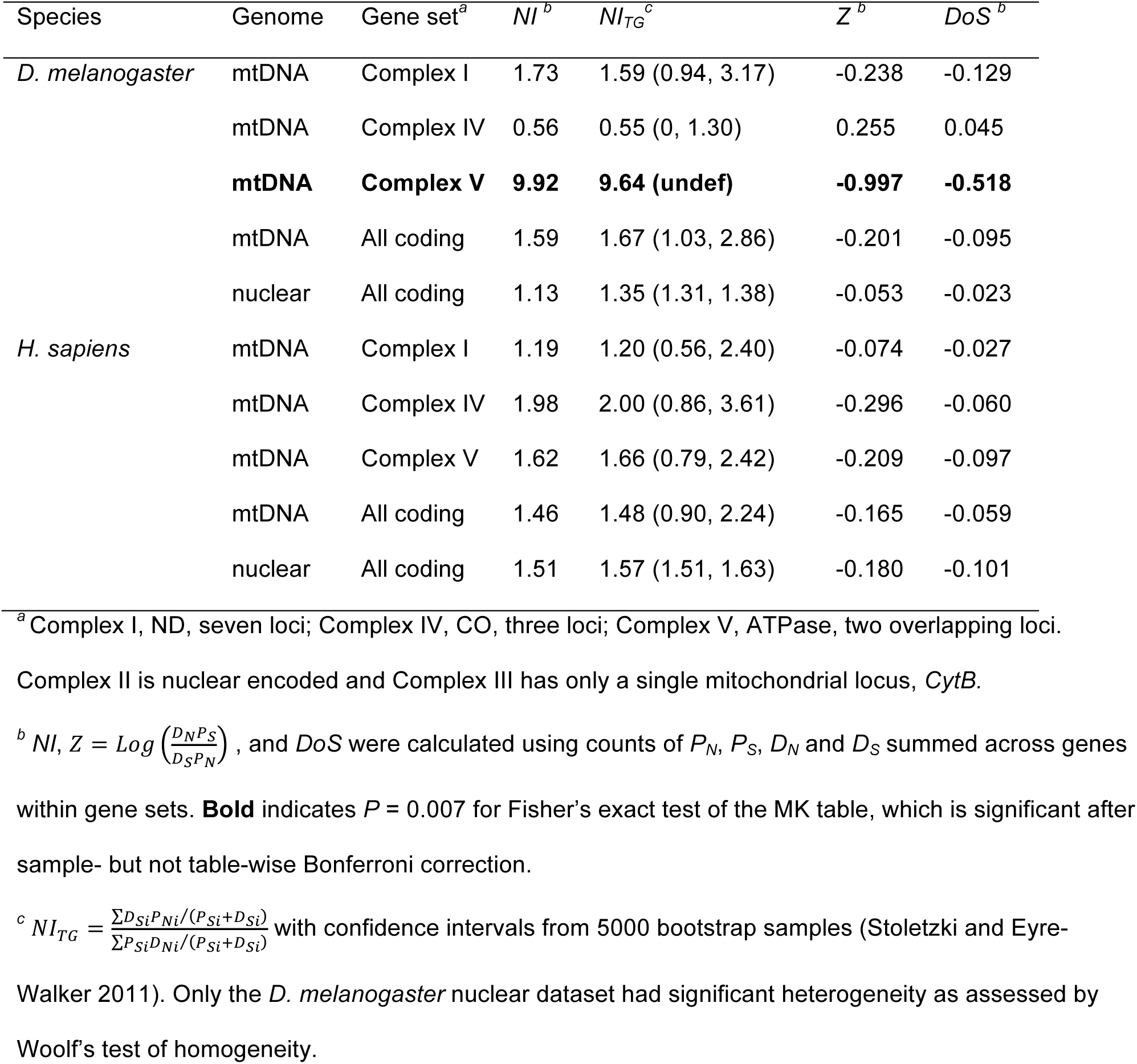
Summary statistics of the McDonald-Kreitman table for mitochondrial OXPHOS complexes and nuclear genes.

| Species | Genome | Gene set<sup>a</sup> | $NI^b$ | $NI_{TG}^c$ | $Z^b$ | $DoS^b$ |
| --- | --- | --- | --- | --- | --- | --- |
| *D. melanogaster* | mtDNA | Complex I | 1.73 | 1.59 (0.94, 3.17) | -0.238 | -0.129 |
|  | mtDNA | Complex IV | 0.56 | 0.55 (0, 1.30) | 0.255 | 0.045 |
|  | **mtDNA** | **Complex V** | **9.92** | **9.64 (undef)** | **-0.997** | **-0.518** |
|  | mtDNA | All coding | 1.59 | 1.67 (1.03, 2.86) | -0.201 | -0.095 |
|  | nuclear | All coding | 1.13 | 1.35 (1.31, 1.38) | -0.053 | -0.023 |
| *H. sapiens* | mtDNA | Complex I | 1.19 | 1.20 (0.56, 2.40) | -0.074 | -0.027 |
|  | mtDNA | Complex IV | 1.98 | 2.00 (0.86, 3.61) | -0.296 | -0.060 |
|  | mtDNA | Complex V | 1.62 | 1.66 (0.79, 2.42) | -0.209 | -0.097 |
|  | mtDNA | All coding | 1.46 | 1.48 (0.90, 2.24) | -0.165 | -0.059 |
|  | nuclear | All coding | 1.51 | 1.57 (1.51, 1.63) | -0.180 | -0.101 |

<sup>a</sup> Complex I, ND, seven loci; Complex IV, CO, three loci; Complex V, ATPase, two overlapping loci.
Complex II is nuclear encoded and Complex III has only a single mitochondrial locus, *CytB*.
<sup>b</sup> $NI$ , $Z = \text{Log} \left( \frac{D_N P_S}{D_S P_N} \right)$ , and $DoS$ were calculated using counts of $P_N$ , $P_S$ , $D_N$ and $D_S$ summed across genes within gene sets. **Bold** indicates $P = 0.007$ for Fisher's exact test of the MK table, which is significant after sample- but not table-wise Bonferroni correction.
<sup>c</sup> $NI_{TG} = \frac{\sum D_{Si} P_{Ni} / (P_{Si} + D_{Si})}{\sum P_{Si} D_{Ni} / (P_{Si} + D_{Si})}$ with confidence intervals from 5000 bootstrap samples (Stoletzki and Eyre-Walker 2011). Only the *D. melanogaster* nuclear dataset had significant heterogeneity as assessed by Woolf's test of homogeneity.

These patterns were verified using 36 of the 38 mitochondrial haplotypes in our sample that were independently sequenced and assembled by Richardson et al. (2012). No single mitochondrial gene rejected the neutral expectation and the MK table counts for Complex V (ATPase) were exactly the same in the two data sets (Table S3). When counts of polymorphism and divergence differed between our datasets, they typically differed by only a single count. The only exception was in several Complex I (ND) genes, for which our assembled mtDNAs had a small number of additional nonsynonymous polymorphisms relative to the Richardson et al. (2012) data set (ND genes, *df* = 6, *P*_*MWU, paired*_ = 0.021; all other genes, *df* = 5, *P*_*MWU, paired*_ = 1) that resulted in slightly higher values of *NI* (ND included, *df* = 12, *P*_*MWU, paired*_ = 0.016; all other genes, *df* = 5, *P*_*MWU, paired*_ = 1). This was not due to exclusion of two mitochondrial haplotypes in the Richardson et al. (2012) data set, and the additional polymorphisms in our data were not clustered on any single haplotype. The reduced number of nonsynonymous polymorphisms in the Richardson et al. (2012) data provided even less support for an excess of nonsynonymous segregating variation in the mitochondrial genome. Patterns of polymorphism and divergence for the entire set of mitochondrial-encoded proteins in this dataset did not deviate from the neutral expectation (*P*_*FET*_ = 0.423), and the confidence intervals on *NI*_*TG*_ for mitochondrially encoded proteins contained the neutral expectation of 1 (*NI*_*TG*_ = 0.821, 95% CI = 0.386 to 1.90).

### On average, *NI* is the same for mitochondrial and nuclear genes in flies

The distribution of *D. melanogaster* mitochondrial gene *NI* values was contained within that of the 6,151 nuclear genes for which we had MK table counts, with many nuclear genes having both more positive and more negative values of *Z\** and *DoS* (Figure 1 and Figure 4). The median value of *NI* was 1.97 (*NI*_*TG*_ = 1.67) for the 13 mitochondrial genes and 1.48 (*NI*_*TG*_ = 1.35) for 6,113 nuclear genes, and did not differ significantly between genomes (*P*_*MWU*_ = 0.278). Confidence intervals for *NI*_*TG*_ were overlapping for the two genomes (Table 3). Median values of *Z\** and *DoS* were negative for both genomes (Figure 1 and Figure 4), consistent with the presence of a slight excess of deleterious polymorphisms in both genomes. *Z\** for *D. melanogaster* mitochondrial genes did not differ significantly from nuclear genes, independent of nuclear location (mtDNA vs. X chromosome, *P*_*MWU*_ = 0.065; mtDNA vs. autosomes, *P*_*MWU*_ = 0.325) (Figure 1C). *DoS* was modestly significantly different between mitochondrial and X-chromosome genes (mtDNA vs. X, *P*_*MWU*_ = 0.044), but not between the mtDNA and autosomal genes (mtDNA vs. autosomes, *P*_*MWU*_ = 0.126). Nevertheless, there was a trend for mitochondrial genes to differ more from X-linked loci than they did from autosomal loci for both *Z\** and *DoS* (Figure 1C,D). Analysis of assembled mtDNAs from Richardson et al. (2012) (see above) yielded qualitatively similar results; *NI* did not differ between mitochondrial and nuclear genes (*P*_*MWU*_ = 0.500), and neither *Z\** or *DoS* differed significantly between the mitochondria and any nuclear location (*Z\**, *P*_*MWU*_ > 0.471 for both comparisons; *DoS*, *P*_*MWU*_ > 0.726 for both comparisons).

**Figure 4.**
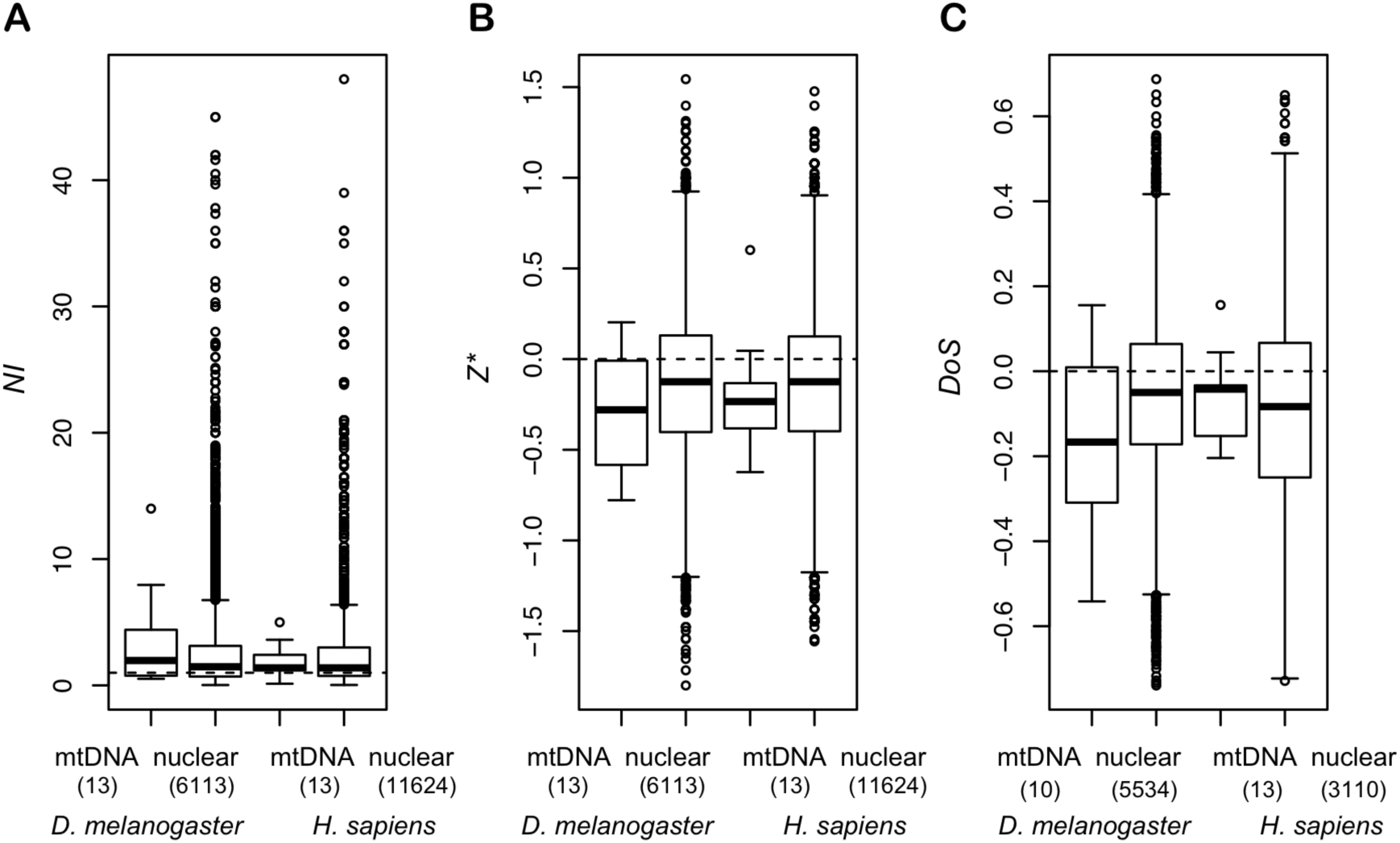
Distributions of polymorphism and divergence summary statistics for mitochondrial and nuclear genes in flies and in humans. (A) Distributions of *NI* are similar between genomes within species and across species. Values of *NI* greater than 50 were removed to improve visualization (3 nuclear genes for flies and 2 nuclear genes for humans). Distributions of (B) *Z\** and (C) *DoS* for mitochondrial genes are also similar to those for nuclear genes in both flies and humans (see text for statistical contrasts). Positive values of these summary statistics of the MK contingency table are consistent with positive selection and negative values with negative selection. Dashed lines represent the neutral expectation for each statistic, and the numbers of genes in each set are indicated in parentheses.

### *NI* does not differ between mitochondrial and nuclear genes in humans

Summary statistics of the MK table also did not differ between mitochondrial and nuclear genes in *H. sapiens* (*NI, P_MWU_* = 0.657; *Z\**, *P*_*MWU*_ = 0.243; *DoS, P_MWU_* = 0.700)(Figure 4), nor did the site-frequency spectrum differ between non-synonymous and synonymous mitochondrial polymorphisms (heterogeneity test, *P* > 0.36 for all genes). Values of *Z\** and *DoS* for mitochondrial genes in humans were well within the distribution of the 11,624 nuclear genes examined, and the confidence intervals of *NI*_*TG*_ were overlapping for the mitochondrial and nuclear genomes (Table 3). The distributions of mitochondrial gene MK summary statistics were also largely overlapping and did not differ significantly between *D. melanogaster* and *H. sapiens* (*NI, P_MWU_* = 0.545; *Z\**, *P*_*MWU*_ = 0.441; *DoS, P_MWU_* = 0.310)(Figure 4), despite differing nuclear *N*_*e*_ between these species. This further supports the idea that the efficacy of selection in the mitochondrial genome is largely independent of *N*_*e*_.

We also found that *NI* is very sensitive to sampling (Table 4). When only a few individuals are sampled, the choice of genomes can lead to high variability and extreme values in *NI*, potentially as a result of single haplotypes that may carry multiple polymorphisms, as appears to be the case for human *ND6* (Table 4). For example, depending on which Japanese individual we included in our analyses, *NI* for *ND6* takes on values of 30.71 (MK, *P*_*FET*_ = 0.001), 5.50 (MK, *P*_*FET*_ = 0.31), or 1.79 (MK, *P*_*FET*_ = 0.16) when sampling only 3 mtDNAs. As more mtDNAs are sampled, point estimates of *NI* and *Z\** for each mitochondrial gene become more similar to the neutral expectation (Table 4) and average values of *NI* and *NI*_*TG*_ no longer show strong evidence for deviations from the neutral expectation.

**Table 4.**
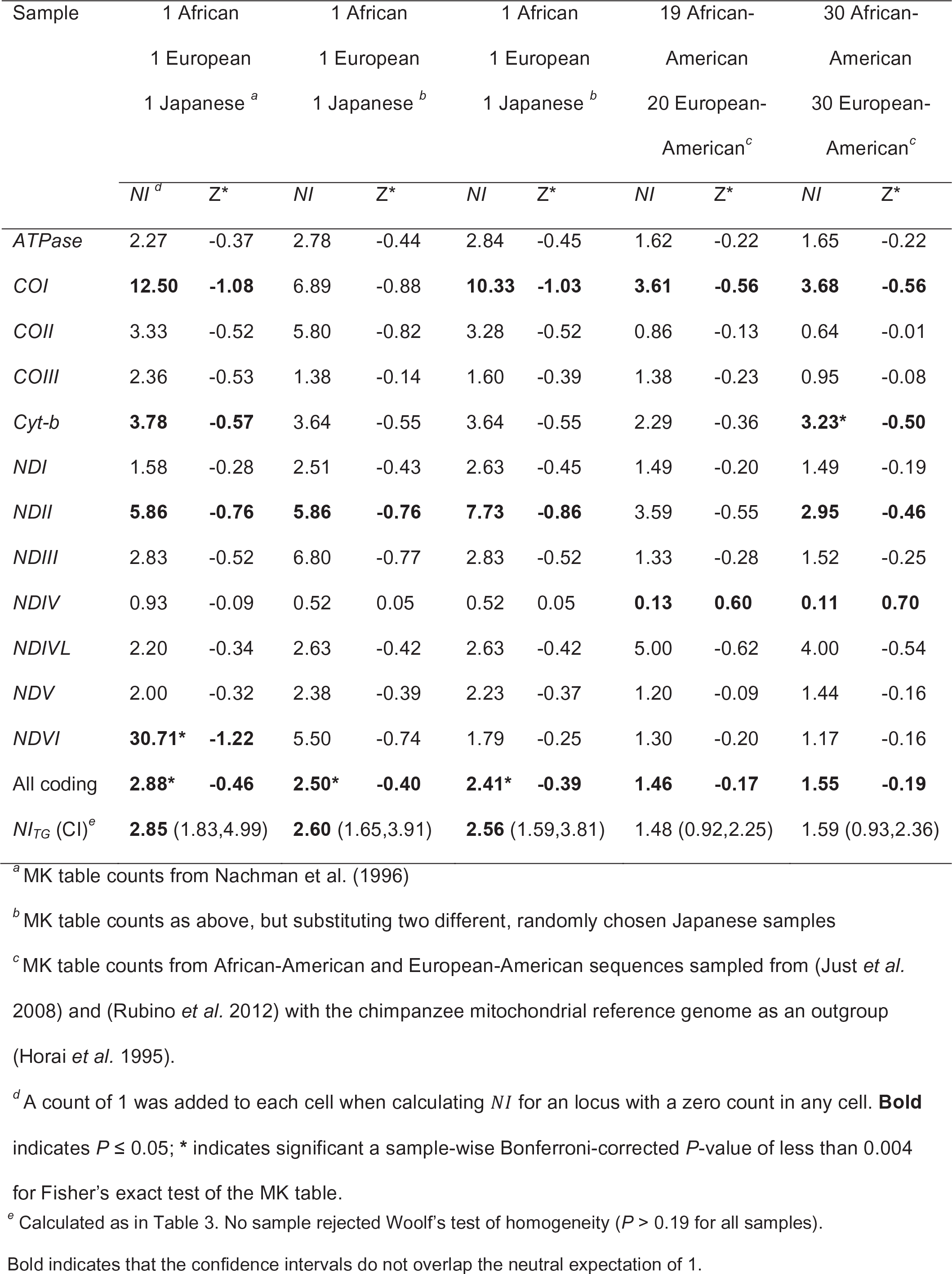
The sensitivity of *NI* to sampling.

## DISCUSSION

Using a large sample of whole-genome sequence data, we have tested a number of hypotheses about mtDNA evolution, and about differences in the efficacy of selection on mitochondrial versus nuclear genes. Our data confirm that mtDNA do not recombine and have lower silent site diversity than do nuclear genes, which supports the prediction that the mitochondrial genome has a lower *N*_*e*_ than does the nuclear genome. We also show a skew in the site-frequency spectrum toward rare alleles that likely has two sources: 1) the accumulation of new mutations on what appears to be a mtDNA haplotype that has swept to high frequency in the recent past, and 2) the ancestral polymorphisms contained on migrant or remnant haplotypes (RAL-639 and RAL-335) that are now rare in this population. Despite the apparent reduction in *N*_*e*_ for mtDNA, our findings indicate that selection is equally effective at purging deleterious polymorphisms from the mitochondrial and nuclear genomes of *D. melanogaster* and *H. sapiens*.

Given its uniparental and haploid transmission, the expectation under neutrality is that the mtDNA has one-quarter the population size of the autosomes. This reduced value of *N* (and subsequently *N*_*e*_) matches that expected for the Y (or W) chromosome, and, like the Y, the mtDNA has little to no recombination. However, very much unlike the Y chromosomes that have been sequenced (e.g. Charlesworth and Charlesworth 2000; Carvalho *et al.* 2009; Carvalho and Clark 2013; Bellott *et al.* 2014), animal mtDNA genomes do not show an accumulation of transposable elements, and the gene content of the mitochondrial genome is remarkably stable, with few gene losses and even fewer pseudogenes (Boore 1999; Ballard and Rand 2005). Furthermore, *d*_*N*_/*d*_*S*_ is two to fifteen times lower for mitochondrial genes than for nuclear genes in mammals (Popadin *et al.* 2012), and average values of *d*_*N*_/*d*_*S*_ for mitochondrial genes are well under 0.1 and are, on average, only 13% that of nuclear genes in *Drosophila* (Bazin *et al.* 2006; Montooth *et al.* 2009). This pattern of amino acid conservation is particularly striking given that the mutation rate in the *D. melanogaster* mtDNA is an order of magnitude greater than the per-site mutation rate in the nuclear genome (Haag-Liautard *et al.* 2007; Haag-Liautard *et al.* 2008). While heteromorphic Y chromosomes do show signatures of less effective purifying selection, such as proliferation of satellite repeats and reduced codon bias (Bachtrog 2013; Singh *et al.* 2014), the single copy, X-degenerate genes that have remained on the human Y chromosome experience effective purifying selection (Rozen *et al.* 2009; Bellott *et al.* 2014), as do the protein sequences of *Drosophila* Y-linked genes (Singh *et al.* 2014). Thus, despite early loss of many genes when heteromorphic Y chromosomes and mtDNA formed, both these non-recombining chromosomes have maintained a stable set of genes that experience effective purifying selection despite their reduced *N*_*e*_.

Many researchers have cited the early work on *NI* in *Drosophila* and mammals in support of the idea that mtDNA accumulate deleterious mutations (e.g., Meiklejohn *et al.* 2007; Green *et al.* 2008; Neiman and Taylor 2009; Akashi *et al.* 2012). In fact, this idea has become so engrained that it is regularly cited in reviews of mitochondrial gene evolution (e.g. Ballard and Whitlock 2004; Lynch 2007, p. 338). What is surprising about this conversion of a small set of intriguing initial studies into dogma is that the early studies themselves were quite circumspect about the implications of their results. For instance, Nachman (1998), in noting that very few nuclear loci were available for comparison, stated “It is also unclear whether the patterns reported here are unique to mitochondrial DNA.” Data from the few nuclear genes that had been sequenced raised “the possibility that the patterns reported here for mtDNA may also be found at some nuclear loci” (Nachman 1998). Even the relatively more recent studies that did have access to additional nuclear datasets were only able to calculate *NI* for 36 nuclear loci (Weinreich and Rand 2000), and the *NI* values that were available often did not deviate significantly from neutrality (Nachman 1998; Weinreich and Rand 2000). Those that did reject neutrality tended to do so weakly, perhaps due to the small number of polymorphisms in mitochondrial samples even when the number of individuals sampled is high (e.g. ND3, Nachman *et al.* 1996). Nevertheless, there are mitochondrial genes that do strongly reject neutrality and some of these had *NI* values that greatly exceeded *NI* for the sampled nuclear loci. Based on these and similar comparisons, many authors have reached the conclusion that mtDNA evolves in a manner distinct from the nuclear genome. Our results using the whole genomes of flies and humans suggest that they are not evolving differently.

Reductions in *N*_*e*_—due either to reductions in census population size or to the increased effect of linked selected variants in regions of low recombination—are expected to result in a reduction in the efficacy of selection. Indeed, comparisons of McDonald-Kreitman test results across a range of species with different values of *N*_*e*_ has revealed this expected relationship (e.g. Li *et al.* 2008; Wright and Andolfatto 2008; Gossmann *et al.* 2010), as have comparisons of *NI* across regions of the *D. melanogaster* genome with different recombination rates (Presgraves 2005; Langley *et al.* 2012). Therefore, all things being equal, mitochondrial loci would be expected to harbor an excess of replacement polymorphisms relative to nuclear loci due to reduced *N*_*e*_. Our results suggest that all things are not equal between these two cellular compartments, and that there may be features of the mitochondrion that make it less likely to accumulate deleterious mutations. One such feature is the “bottleneck” that occurs in the number of mtDNAs that are passed from mother to offspring—this event makes it possible for selection to act within hosts, possibly increasing the power of selection to remove deleterious mutations (Bergstrom and Pritchard 1998; Rand 2011) and reducing variability in mitochondrial *N*_*e*_ among taxa, relative to nuclear genomes. The additional layers of selection imposed by mitochondrial inheritance, combined with stronger negative selective effects of amino acid changing mutations in mitochondrial genes, may therefore allow the mtDNA to escape the accumulation of deleterious mutation, resulting in equal values of *NI* between nucleus and mitochondria. If the selective effects of mutations in mitochondrial genes are beyond the “horizons” where all mutations will behave similarly regardless of *N*_*e*_ (Nachman 1998; Eyre-Walker and Keightley 2007), then we expect patterns of mitochondrial polymorphism and divergence to be largely independent of *N*_*e*_.

Our results come with several caveats. First, we have only studied two organisms—it may be that a more comprehensive review of *NI* in mtDNA and nuclear loci across many species will reveal a difference in the average efficacy of selection. The early meta-analyses of *NI* contained loci from a wide range of animals (Nachman 1998; Weinreich and Rand 2000), possibly suggesting that using only *Drosophila* and humans results in a view that is too limited. Nevertheless, these are two model organisms for evolutionary biology that span a large range of mtDNA:nuclear substitution rates, and studies of these species have set the bar for much of modern population genetics. Second, it is clear from our analysis of the *D. melanogaster* mtDNA that it is not at equilibrium, and may be recovering from a partial cytoplasmic sweep that may be associated with *Wolbachia* (Richardson et al. 2012). Much of the theory used to predict *NI* values from *N*_*e*_ and *s* assumes mutation-selection-drift balance (see, e.g. Nachman 1998), and deviations from this equilibrium can result in more complex relationships between *N*_*e*_, *s*, and *NI* (Messer and Petrov 2013). While non-equilibrium histories may mean that mtDNA *NI* values are not at equilibrium, it is equally likely that nuclear genes from *D. melanogaster* are not at mutation-selection-drift equilibrium (Hahn 2008; Langley *et al.* 2012). Whether or not the mtDNA is at equilibrium, and whether or not the *NI* values calculated from this snapshot of two species represent equilibrium values, our results still imply that there is no difference between nuclear and mitochondrial measures of the efficacy of selection.

Despite the mitochondrial genome experiencing a distinct population genetic environment relative to the nuclear genome, our whole-genome analyses uncovered no evidence for an excess accumulation of slightly deleterious mutations in mitochondrial genomes. In fact, the only strong evidence for a reduced efficacy of selection in mtDNA, relative to nuclear genomes, comes from comparative studies of nuclear and mitochondrial tRNAs (Lynch 1996; Lynch 1997). As discussed above, in the absence of a pattern in *NI*, there are essentially no other patterns of molecular evolution in mtDNA indicative of deleterious mutation accumulation. This pattern is in stark contrast to the patterns found in analogous nuclear regions with reduced *N*_*e*_ and low recombination, like the Y chromosome. Determining whether mtDNA accumulate deleterious polymorphisms and substitutions more readily than nuclear DNA in a larger sample of species—and what type of loci may be affected—will be a particularly fruitful avenue for future studies.

## ACKNOWLEDGEMENTS

We thank Colin Meiklejohn and members of the Montooth and Hahn labs for constructive feedback. B.S.C. was supported on the Indiana University Genetics, Molecular and Cellular Sciences Training Grant T32-GM007757 funded by the National Institutes of Health and a Doctoral Dissertation Improvement Grant funded by the National Science Foundation. This research was supported by funding from Indiana University, award DBI-0845494 from the National Science Foundation to M.W.H., and a National Science Foundation CAREER award IOS-1149178 to K.L.M.

## Supporting Information

**File S1** FASTA file with the 38 assembled and aligned genomes used in this study

**Table S1** mtDNAs assembled in this study along with the average coverage

**Table S2** Correlations between linkage disequilibrium (LD) and distance (bp) between pairs of SNPs in the *D. melanogaster* mtDNA

**Table S3** Counts of polymorphic (*P*) and divergent (*D*) nonsynonymous (*N*) and synonymous (*S*) sites along with summary statistics of the McDonald-Kreitman table using 36 of the 38 mitochondrial haplotypes in our sample that were independently sequenced and assembled by Richardson et al. (2012).

**Table S4** Effects of sampling on *NI*

**Table S5** Accession numbers for mtDNAs used in Table S4

## LITERATURE CITED

Akashi, H., N. Osada and T. Ohta, 2012 Weak selection and protein evolution. Genetics 192: 15–470 U43.

Awadalla, P., A. Eyre-Walker and J. M. Smith, 1999 Linkage disequilibrium and recombination in hominid mitochondrial DNA. Science 286: 2524–2525.

Bachtrog, D., 2013 Y-chromosome evolution: emerging insights into processes of Y-chromosome degeneration. Nature Reviews Genetics 14: 113–124.

Ballard, J. W., 2000 Comparative genomics of mitochondrial DNA in members of the *Drosophila melanogaster* subgroup. J Mol Evol 51: 48–63.

Ballard, J. W., and M. Kreitman, 1994 Unraveling selection in the mitochondrial genome of *Drosophila*. Genetics 138: 757–772.

Ballard, J. W. O., and D. M. Rand, 2005 The population biology of mitochondrial DNA and its phylogenetic implications, pp. 621–642 in Annu Rev Ecol Evol S.

Ballard, J. W. O., and M. C. Whitlock, 2004 The incomplete natural history of mitochondria. Molecular Ecology 13: 729–744.

Bazin, E., S. Glemin and N. Galtier, 2006 Population size does not influence mitochondrial genetic diversity in animals. Science 312: 570–572.

Bellott, D. W., J. F. Hughes, H. Skaletsky, L. G. Brown, T. Pyntikova et al., 2014 Mammalian Y chromosomes retain widely expressed dosage-sensitive regulators. Nature 508: 494-+.

Bergstrom, C. T., and J. Pritchard, 1998 Germline bottlenecks and the evolutionary maintenance of mitochondrial genomes. Genetics 149: 2135–2146.

Betancourt, A. J., B. Blanco-Martin and B. Charlesworth, 2012 The relation between the neutrality index for mitochondrial genes and the distribution of mutational effects on fitness. Evolution 66: 2427–2438.

Boore, J. L., 1999 Animal mitochondrial genomes. Nucleic Acids Res 27: 1767–1780.

Bruen, T. C., H. Philippe and D. Bryant, 2006 A simple and robust statistical test for detecting the presence of recombination. Genetics 172: 2665–2681.

Bustamante, C. D., A. Fledel-Alon, S. Williamson, R. Nielsen, M. T. Hubisz et al., 2005 Natural selection on protein-coding genes in the human genome. Nature 437: 1153–1157.

Carvalho, A. B., and A. G. Clark, 2013 Efficient identification of Y chromosome sequences in the human and *Drosophila* genomes. Genome Research 23: 1894–1907.

Carvalho, A. B., L. B. Koerich and A. G. Clark, 2009 Origin and evolution of Y chromosomes: *Drosophila* tales. Trends in Genetics 25: 270–277.

Charlesworth, B., 2012 The effects of deleterious mutations on evolution at linked sites. Genetics 190: 5–22.

Charlesworth, B., and D. Charlesworth, 2000 The degeneration of Y chromosomes. Philosophical Transactions of the Royal Society of London. Series B: Biological Sciences 505 355: 1563–1572.

Clary, D. O., and D. R. Wolstenholme, 1985 The mitochondrial DNA molecular of *Drosophila yakuba*: nucleotide sequence, gene organization, and genetic code. J Mol Evol 22: 252–271.

Clement, M., D. Posada and K. A. Crandall, 2000 TCS: a computer program to estimate gene genealogies. Mol Ecol 9: 1657-1659.

Eyre-Walker, A., and P. D. Keightley, 2007 The distribution of fitness effects of new mutations. Nat Rev Genet 8: 610–618.

Fay, J. C., and C. I. Wu, 2000 Hitchhiking under positive Darwinian selection. Genetics 155: 1405–1413.

Fu, Y. X., and W. H. Li, 1993 Statistical tests of neutrality of mutations. Genetics 133: 693–709.

Gillespie, J. H., 2000 Genetic drift in an infinite population: The pseudohitchhiking model. Genetics 155: 909–919.

Gossmann, T. I., B.-H. Song, A. J. Windsor, T. Mitchell-Olds, C. J. Dixon et al., 2010 Genome wide qnalyses reveal little evidence for adaptive evolution in many plant species. Molecular Biology and Evolution 27: 1822–1832.

Green, R. E., A.-S. Malaspinas, J. Krause, A. W. Briggs, P. L. F. Johnson et al., 2008 A complete neandertal mitochondrial genome sequence determined by high-throughput Sequencing. Cell 134: 416–426.

Haag-Liautard, C., N. Coffey, D. Houle, M. Lynch, B. Charlesworth et al., 2008 Direct estimation of the mitochondrial DNA mutation rate in *Drosophila melanogaster*. PLoS Biol 525 6: e204.

Haag-Liautard, C., M. Dorris, X. Maside, S. Macaskill, D. L. Halligan et al., 2007 Direct estimation of per nucleotide and genomic deleterious mutation rates in *Drosophila*. Nature 445: 82–85.

Hahn, M. W., 2008 Toward a selection theory of molecular evolution. Evolution 62: 255–265.

Hahn, M. W., M. D. Rausher and C. W. Cunningham, 2002 Distinguishing between selection and population expansion in an experimental lineage of bacteriophage t7. Genetics 161: 11–20.

Hill, W. G., and A. Robertson, 1966 The effect of linkage on limits to artificial selection. Genet Res 8: 269–294.

Horai, S., K. Hayasaka, R. Kondo, K. Tsugane and N. Takahata, 1995 Recent African origin of modern humans revealed by complete sequences of hominoid mitochondrial DNAs. Proc Natl Acad Sci U S A 92: 532–536.

Hudson, R. R., and N. L. Kaplan, 1985 Statistical properties of the number of recombination events in the history of a sample of DNA sequences. Genetics 111: 147–164.

Innan, H., and M. Nordborg, 2002 Recombination or mutational hot spots in human mtDNă Mol Biol Evol 19: 1122–1127.

Just, R. S., T. M. Diegoli, J. L. Saunier, J. A. Irwin and T. J. Parsons, 2008 Complete mitochondrial genome sequences for 265 African American and U.S. “Hispanic” individuals. Forensic Sci Int Genet 2: e45–48.

Kimura, M., 1983 The neutral theory of molecular evolution. Cambridge University Press, Cambridge.

Langley, C., K. Stevens, C. Cardeno, Y. Lee, D. Schrider et al., 2012 Genomic variation in natural populations of *Drosophila melanogaster*. Genetics 192: 533–598.

Larkin, M. A., G. Blackshields, N. P. Brown, R. Chenna, P. A. Mcgettigan et al., 2007 Clustal W and clustal X version 2.0. Bioinformatics 23: 2947–2948.

Lewontin, R. C., 1964 The interaction of selection and linkage. I. General considerations; heterotic models. Genetics 49: 49–67.

Li, H., and R. Durbin, 2009 Fast and accurate short read alignment with Burrows-Wheeler transform. Bioinformatics 25: 1754–1760.

Li, H., B. Handsaker, A. Wysoker, T. Fennell, J. Ruan et al., 2009 The Sequence Alignment/Map format and SAMtools. Bioinformatics 25: 2078–2079.

Li, Y. F., J. C. Costello, A. K. Holloway and M. W. Hahn, 2008 “Reverse ecology” and the power of population genomics. Evolution 62: 2984–2994.

Lynch, M., 1996 Mutation accumulation in transfer RNAs: molecular evidence for Muller's ratchet in mitochondrial genomes. Mol Biol Evol 13: 209–220.

Lynch, M., 1997 Mutation accumulation in nuclear, organelle, and prokaryotic transfer RNA genes. Mol Biol Evol 14: 914–925.

Lynch, M., 2007 Origins of Genome Architecture. Sinauer Associates, Inc., Sunderland, Massachusetts.

Lynch, M., W. Sung, K. Morris, N. Coffey, C. R. Landry et al., 2008 A genome-wide view of the spectrum of spontaneous mutations in yeast. Proc Natl Acad Sci USA.

Mackay, T., S. Richards, E. Stone, A. Barbadilla, J. Ayroles et al., 2012 The *Drosophila melanogaster* genetic reference panel. Nature 482: 173–178.

Maynard Smith, J., and J. Haigh, 1974 The hitch-hiking effect of a favourable gene. Genetical Research 23: 23–35.

Mcdonald, J. H., and M. Kreitman, 1991 Adaptive protein evolution at the *Adh* locus in *Drosophila*. Nature 351: 652–654.

Meiklejohn, C. D., K. L. Montooth and D. M. Rand, 2007 Positive and negative selection on the mitochondrial genome. Trends Genet 23: 259–263.

Messer, P. W., and D. A. Petrov, 2013 Frequent adaptation and the McDonald–Kreitman test. Proceedings of the National Academy of Sciences 110: 8615–8620.

Meunier, J., and A. Eyre-Walker, 2001 The correlation between linkage disequilibrium and distance: implications for recombination in hominid mitochondria. Mol Biol Evol 18: 2132–2135.

Montooth, K. L., D. N. Abt, J. Hofmann and D. M. Rand, 2009 Comparative genomics of *Drosophila* mtDNA: Novel features of conservation and change across functional domains and lineages. J Mol Evol 69: 94–114.

Montooth, K. L., and D. M. Rand, 2008 The spectrum of mitochondrial mutation differs across species. PLoS Biol 6: e213.

Nachman, M. W., 1998 Deleterious mutations in animal mitochondrial DNA. Genetica 102-103: 61–69.

Nachman, M. W., W. M. Brown, M. Stoneking and C. F. Aquadro, 1996 Nonneutral mitochondrial DNA variation in humans and chimpanzees. Genetics 142: 953–963.

Nei, M., and T. Gojobori, 1986 Simple methods for estimating the numbers of synonymous and nonsynonymous nucleotide substitutions. Mol Biol Evol 3: 418–426.

Neiman, M., and D. R. Taylor, 2009 The causes of mutation accumulation in mitochondrial genomes. Proc Biol Sci 276: 1201–1209.

Popadin, K. Y., S. I. Nikolaev, T. Junier, M. Baranova and S. E. Antonarakis, 2012 Purifying selection in mammalian mitochondrial protein-coding genes is highly effective and congruent with evolution of nuclear genes. Mol Biol Evol.

Presgraves, D., 2005 Recombination enhances protein adaptation in *Drosophila melanogaster*. Curr Biol 15: 1651–1656.

R Core Team, 2012 R: A language and environment for statistical computing. R Foundation for Statistical Computing, Vienna, Austria ISBN 3-900051-07-0: URL http://www.R-project.org/.

Rand, D. M., 2011 Population genetics of the cytoplasm and the units of selection on mitochondrial DNA in *Drosophila melanogaster*. Genetica 139: 685–697.

Rand, D. M., and L. M. Kann, 1996 Excess amino acid polymorphism in mitochondrial DNA: contrasts among genes from *Drosophila*, mice, and humans. Mol Biol Evol 13: 735–748.

Rand, D. M., and L. M. Kann, 1998 Mutation and selection at silent and replacement sites in the evolution of animal mitochondrial DNA. Genetica 102-103: 393–407.

Richardson, M. F., L. A. Weinert, J. J. Welch, R. S. Linheiro, M. M. Magwire et al., 2012 Population genomics of the *Wolbachia* endosymbiont in *Drosophila melanogaster*. PLoS genetics 8: e1003129.

Rozas, J., J. C. Sanchez-Delbarrio, X. Messeguer and R. Rozas, 2003 DnaSP, DNA polymorphism analyses by the coalescent and other methods. Bioinformatics 19: 2496–2497.

Rozen, S., J. D. Marszalek, R. K. Alagappan, H. Skaletsky and D. C. Page, 2009 Remarkably little variation in proteins encoded by the Y chromosome's single-copy genes, implying effective purifying selection. American Journal of Human Genetics 85: 923–928.

Rubino, F., R. Piredda, F. M. Calabrese, D. Simone, M. Lang et al., 2012 HmtDB, a genomic resource for mitochondrion-based human variability studies. Nucleic Acids Res 40: D1150–1159.

Sheldahl, L. A., D. M. Weinreich and D. M. Rand, 2003 Recombination, dominance and selection on amino acid polymorphism in the *Drosophila* genome: contrasting patterns on the X and fourth chromosomes. Genetics 165: 1195–1208.

Shoemaker, D. D., K. A. Dyer, M. Ahrens, K. Mcabee and J. Jaenike, 2004 Decreased diversity but increased substitution rate in host mtDNA as a consequence of *Wolbachia* endosymbiont infection. Genetics 168: 2049–2058.

Singh, N. D., L. B. Koerich, A. B. Carvalho and A. G. Clark, 2014 Positive and purifying selection on the *Drosophila* Y chromosome. Mol Biol Evol 31: 2612–2623.

Stoletzki, N., and A. Eyre-Walker, 2011 Estimation of the neutrality index. Mol Biol Evol 28: 63–70.

Tajima, F., 1989 Statistical method for testing the neutral mutation hypothesis by DNA polymorphism. Genetics 123: 585–595.

Watterson, G. A., 1975 On the number of segregating sites in genetical models without recombination. Theor Popul Biol 7: 256–276.

Weinreich, D. M., and D. M. Rand, 2000 Contrasting patterns of nonneutral evolution in proteins encoded in nuclear and mitochondrial genomes. Genetics 156: 385–399.

White, D. J., J. N. Wolff, M. Pierson and N. J. Gemmell, 2008 Revealing the hidden complexities of mtDNA inheritance. Mol Ecol 17: 4925–4942.

Wolstenholme, D. R., and D. O. Clary, 1985 Sequence evolution of *Drosophila* mitochondrial DNA. Genetics 109: 725–744.

Woolf, B., 1955 On estimating the relation between blood group and disease. Annals of Human Genetics 19: 251–253.

Wright, S. I., and P. Andolfatto, 2008 The impact of natural selection on the genome: emerging patterns in *Drosophila* and *Arabidopsis*. Annual Review of Ecology Evolution and Systematics 39: 193–213.

